# Identification of novel translated small ORFs in *Escherichia coli* using complementary ribosome profiling approaches

**DOI:** 10.1101/2021.07.02.450978

**Authors:** Anne Stringer, Carol Smith, Kyle Mangano, Joseph T. Wade

## Abstract

Small proteins of <51 amino acids are abundant across all domains of life but are often overlooked because their small size makes them difficult to predict computationally, and they are refractory to standard proteomic approaches. Ribosome profiling has been used to infer the existence of small proteins by detecting the translation of the corresponding open reading frames (ORFs). Detection of translated short ORFs by ribosome profiling can be improved by treating cells with drugs that stall ribosomes at specific codons. Here, we combine the analysis of ribosome profiling data for *Escherichia coli* cells treated with antibiotics that stall ribosomes at either start or stop codons. Thus, we identify ribosome-occupied start and stop codons for ~400 novel putative ORFs with high sensitivity. The newly discovered ORFs are mostly short, with 365 encoding proteins of <51 amino acids. We validate translation of several selected short ORFs, and show that many likely encode unstable proteins. Moreover, we present evidence that most of the newly identified short ORFs are not under purifying selection, suggesting they do not impact cell fitness, although a small subset have the hallmarks of functional ORFs.

**IMPORTANCE:** Small proteins of <51 amino acids are abundant across all domains of life but are often overlooked because their small size makes them difficult to predict computationally, and they are refractory to standard proteomic approaches. Recent studies have discovered small proteins by mapping the location of translating ribosomes on RNA using a technique known as ribosome profiling. Discovery of translated sORFs using ribosome profiling can be improved by treating cells with drugs that trap initiating ribosomes. Here, we show that combining these data with equivalent data for cells treated with a drug that stalls terminating ribosomes facilitates the discovery of small proteins. We use this approach to discover 365 putative genes that encode small proteins in *Escherichia coli*.

## INTRODUCTION

Tens of thousands of bacterial genomes have been fully sequenced. These genome sequences are annotated using computational pipelines that predict the location of open reading frames (ORFs). One of the main criteria used by ORF prediction pipelines is the length of the putative ORFs; ORFs encoding proteins with >50 amino acids (aa) are unlikely to occur by chance; thus, there is a strong bias towards identifying longer ORFs. However, there is strong evidence for large numbers of short ORFs (sORFs; encoding proteins with <50 aa): phylogenetic analysis has identified many conserved sORFs (1), and transcriptomic and proteomic experimental approaches provide strong evidence for the translation of many sORFs (2–15). As more sORFs and their encoded small proteins are identified, it is becoming increasingly clear that sORFs and small proteins play important functional roles in bacteria. In particular, sORFs can function as *cis*-acting regulators of downstream operonic genes, and small proteins can function to modulate the activity of protein complexes (16, 17).

There are 118 described sORFs encoded by *Escherichia coli* K-12, most of which have not been functionally characterized. Recent studies have used genome-scale approaches to identify translated sORFs. Ribosome profiling is a method that experimentally maps ribosome association with the transcriptome. Two studies applied ribosome profiling to *E. coli* BL21 (closely related to *E. coli* K-12) cells treated with antibiotics, retapamulin (“Ribo-RET”) and Onc112, that trap ribosomes at start codons (2, 3). Thus, sites of translation initiation were identified, leading to the identification of many putative translated sORFs. In one study, 38 of the encoded small proteins were successfully validated by western blot (3).

Ribo-RET has also been applied to *Mycobacterium tuberculosis*, revealing an even larger number of putative sORFs than described for *E. coli* (4). Moreover, the majority of the sORFs identified in *M. tuberculosis* do not appear to have been subject to purifying selection, suggesting that they represent the products of “pervasive translation”, whereby ribosomes initiate translation at large numbers of locations across the transcriptome, with many of the translated proteins being non-functional, i.e. not contributing to cell fitness.

Ribosome profiling has also been applied to *E. coli* cells treated with the antibiotic apidaecin, which traps ribosomes at stop codons by binding in the nascent peptide exit tunnel and trapping release factor (18). This method, “Ribo-Api”, also leads to ribosome accumulation upstream and downstream of stop codons, and at start codons (19); this additional signal can be partially reduced by addition of puromycin to cell lysates (“Ribo-Api/Pmn”), but even with this reduction in signal away from stop codons, Ribo-Api/Pmn data cannot be used to identify stop codons with high specificity. Nonetheless, we reasoned that combining Ribo-RET and Ribo-Api/Pmn datasets for *E. coli* would facilitate the identification of translated sORFs with high sensitivity, since the likelihood of detecting Ribo-RET signal at a start codon and Ribo-Api/Pmn signal at the corresponding stop codon by chance would be very low. Our analysis of Ribo-RET and Ribo-Api/Pmn data led to the identification of 397 novel ORFs with high confidence, most of which are sORFs. We validated expression of the associated small proteins for 9 of 18 novel sORFs tested, and we showed that the ribosome-binding sites of all 18 selected sORFs could drive translation of a luciferase reporter assay. Thus, our data suggest that most or all of the 365 putative sORFs identified by combining Ribo-RET and Ribo-Api/Pmn data are robustly translated, but that many of these sORFs encode scarce or unstable small proteins. We speculate that pervasive translation leads to the production of many unstable, non-functional proteins in *E. coli*.

## RESULTS

### Reanalysis of Ribo-RET data to identify putative start codons

Using a previously described Ribo-RET dataset for *E. coli* BL21 (2) (Figure 1A), we identified 12,756 sites of ribosome occupancy that exceed the local background; we refer to these positions as “Initiation-Enriched Ribosome Footprints” (IERFs; Table S1). We then determined the enrichment of every possible trinucleotide sequence in the regions surrounding IERFs. Consistent with most IERFs representing sites of translation initiation, ATG, GTG and TTG trinucleotide sequences were enriched over the local background at positions 15-18 nt upstream of IERFs (Figure 1B), with the highest enrichment observed for ATG sequences, and the lowest enrichment observed for TTG sequences; we did not observe enrichment of CTG, ATT, or ATC sequences, consistent with these sequences being used infrequently as start codons (20). Based on the degree of enrichment for ATG, GTG and TTG sequences at different positions 14-18 nt upstream of IERFs, we reanalyzed the Ribo-RET data to identify putative start codons (see Methods). Thus, we identified 7,936 putative start codons with an estimated false discovery rate (FDR) of 11.7% (see Methods). 2,545 of these putative start codons match those of annotated ORFs. We compared the putative start codons to those predicted previously using the same Ribo-RET data using a higher read coverage threshold, but with a considerably more relaxed range of potential start codons (see methods for details). 116 of the start codons identified in the current study were identified previously in reference (2).

**Figure 1.**
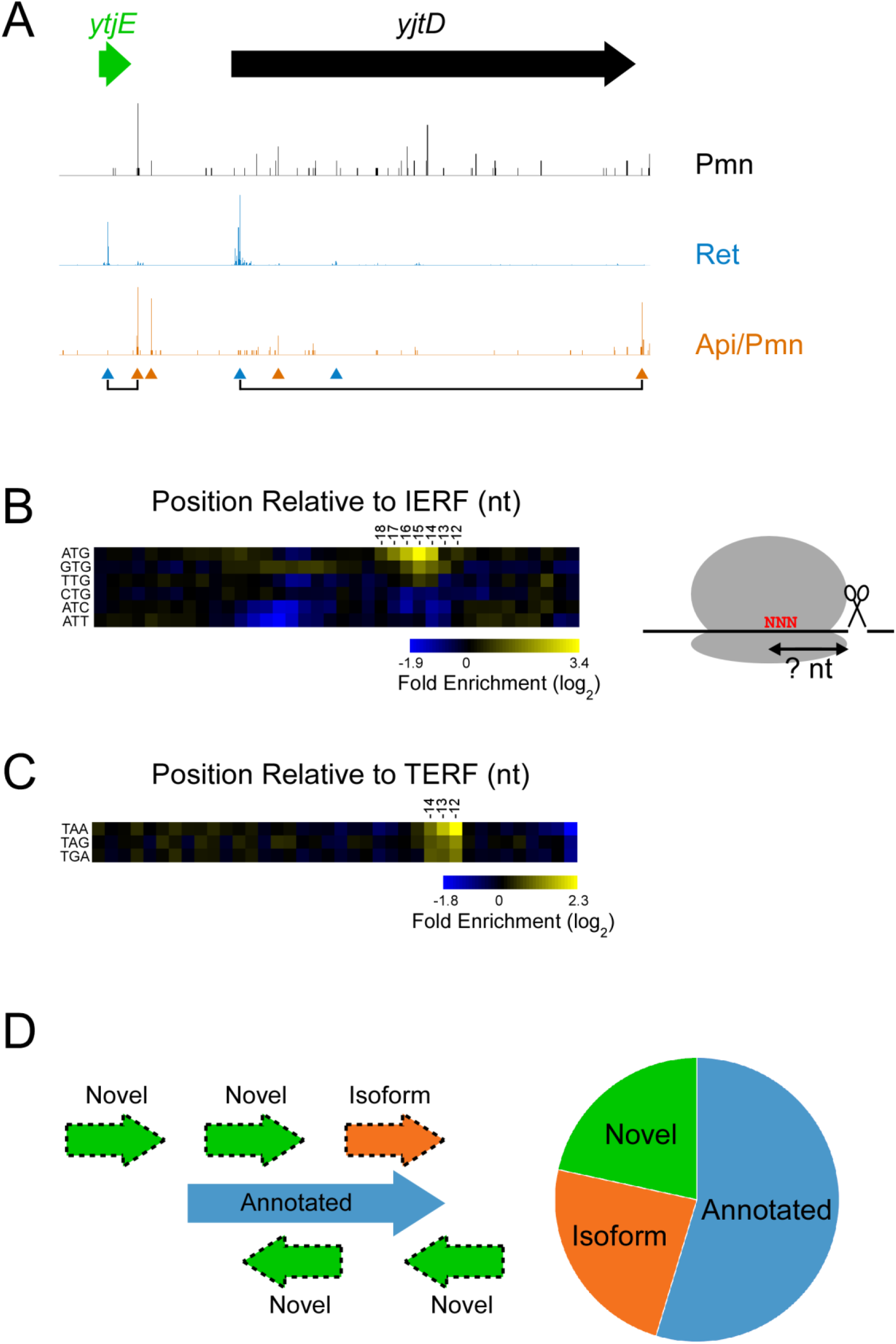
Identification of ORFs by combining Ribo-RET and Ribo-ApiPmn data. **(A)** Sequence read coverage is shown across a selected genomic region for Ribo-RET (retapamulin treatment; stalls initiating ribosomes), Ribo-Api/Pmn (apidaecin and puromycin treatment; stalls terminating ribosomes), and Ribo-Pmn (puromycin treatment; control) data. Blue and orange triangles indicate the positions of start codons paired with Initiation-Enriched Ribosome Footprints (IERFs) and stop codons paired with Termination-Enriched Ribosome Footprints (TERFs), respectively. Triangles joined by lines indicate start and stop codons from the same putative ORF. The positions of the annotated *yjtD* and novel *ytjE* ORFs are shown by horizontal arrows above the data. **(B)** Heatmap showing enrichment of possible start codon sequences at positions upstream of IERFs. **(C)** Heatmap showing enrichment of possible stop codon sequences at positions upstream of TERFs. **(D)** Classification of ORFs identified from Ribo-RET and Ribo-Api/Pmn data, with categories defined based on the overlap of start/stop codons with annotated ORFs, as shown in the schematic.

### Reanalysis of Ribo-Api data to identify putative stop codons

Using a previously described Ribo-Api dataset, including a puromycin treatment (“Ribo-Api/Pmn”), for *E. coli* BL21 (19) (Figure 1A), we identified 12,756 sites of ribosome occupancy that exceed the local background; we refer to these positions as “Termination-Enriched Ribosome Footprints” (TERFs; Table S2). We then determined the enrichment of every possible trinucleotide sequence in the regions surrounding TERFs. Consistent with most TERFs representing sites of translation termination, TAA, TGA and TAA trinucleotide sequences were enriched over the local background at positions 12-14 nt upstream of TERFs (Figure 1C). Based on these enrichments, we reanalyzed the Ribo-Api/Pmn data to identify putative stop codons by selecting TERFs with a TAA/TGA/TAG trinucleotide sequence in a specified distance range upstream. Thus, we identified 6,877 putative stop codons with an estimated FDR of 33.7% (see Methods). 1,377 of these putative stop codons match those of annotated genes.

### Combining putative start and stop codons to identify putative novel ORFs

We reasoned that by selecting putative start codons inferred from Ribo-RET data that are paired with putative stop codons inferred from Ribo-Api/Pmn data, we could identify ORFs with higher confidence than by using Ribo-RET or Ribo-Api/Pmn data alone. Moreover, evidence of ribosome occupancy at both start and stop codons of these putative ORFs implies they are actively translated from beginning to end (Figure 1A). Using this approach, we detected 1,839 putative ORFs (Table S3), with a conservative FDR estimate of 2.2% (see Methods). 1,005 of the ORFs we detected perfectly match annotated genes; we refer to these as “annotated” ORFs. 437 of the ORFs we detected share a stop codon with an annotated gene, but have a different start codon. These could represent mis-annotated genes, or one of a pair of ORFs where the corresponding proteins share the same C-terminal region. We refer to these as “isoform” ORFs. 397 of the ORFs we detected have a stop codon that does not match any annotated gene. We refer to these as “novel” ORFs (Figure 1D). The likelihood of detecting an annotated ORF by chance is exceedingly low (we conservatively estimate the FDR to be ~0.07%; see Methods). We conservatively estimate that the FDRs for identifying isoform and novel ORFs are 2.3% and 7.5%, respectively (see Methods).

### Putative isoform and novel ORF start and stop codons are associated with RNA secondary structure features expected of actively translated ORFs

Actively translated start codons in *E. coli* tend to be associated with regions of reduced RNA secondary structure relative to the surrounding sequence (21). We examined the sequences around the 1,005 annotated, 437 isoform, and 397 novel ORF start codons. For all three classes of ORF, we observed significantly reduced predicted RNA secondary structure in the regions from −25 to +15 nt from start codons, relative to that of 500 randomly selected genomic sequences of the same length (Figure 2; Mann-Whitney U test *p* < 2.2e^-16^ for each of annotated and novel ORFs; *p* = 3.2e^-6^ for isoform ORFs). We conclude that the start codons of isoform and novel ORFs are associated with RNA secondary structure features expected of actively translated ORFs. Predicted RNA secondary structure around the start codons of the annotated ORFs was slightly lower than that for novel ORFs (Mann-Whitney U test *p* = 1.4e^-8^), and substantially lower than that for isoform ORFs (Mann-Whitney U test *p* < 2.2e^-16^).

**Figure 2.**
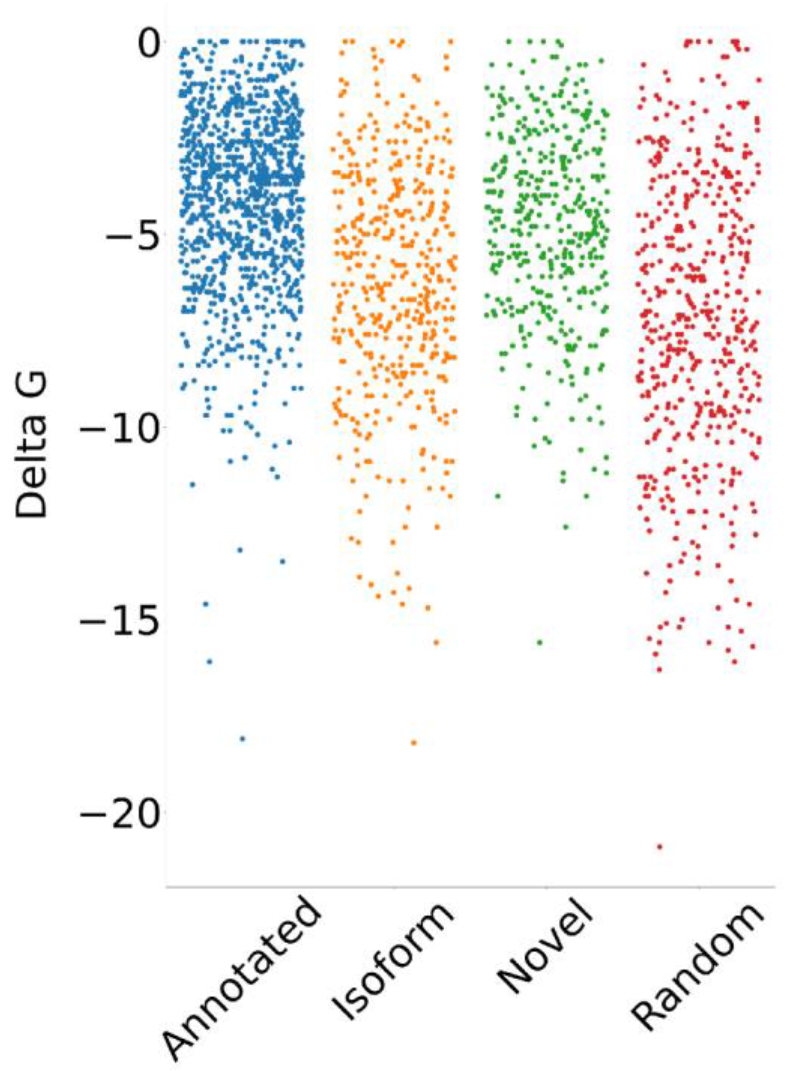
Reduced RNA secondary structure for sequences around identified start codons. Strip plot showing the ΔG for the predicted minimum free energy structures for the regions from −25 to +15 nt relative to putative start codons for annotated, isoform, and novel ORFs identified by analyzing Ribo-RET and Ribo-Api/Pmn data, and for a set of 500 random sequences.

### Classification of novel ORFs based on local gene context

We compared the positions of the 397 putative novel ORFs with the positions of annotated genes (Figure 3). 27% of the novel ORFs are located entirely in intergenic regions; of these, roughly two thirds are immediately upstream of an annotated gene on the same strand, suggesting that they are co-transcribed. 44% of the novel ORFs overlap completely with an annotated gene in the sense orientation, whereas only 12% overlap completely with an annotated gene in the antisense orientation. 17% of the novel ORFs partially overlap an annotated gene.

**Figure 3.**
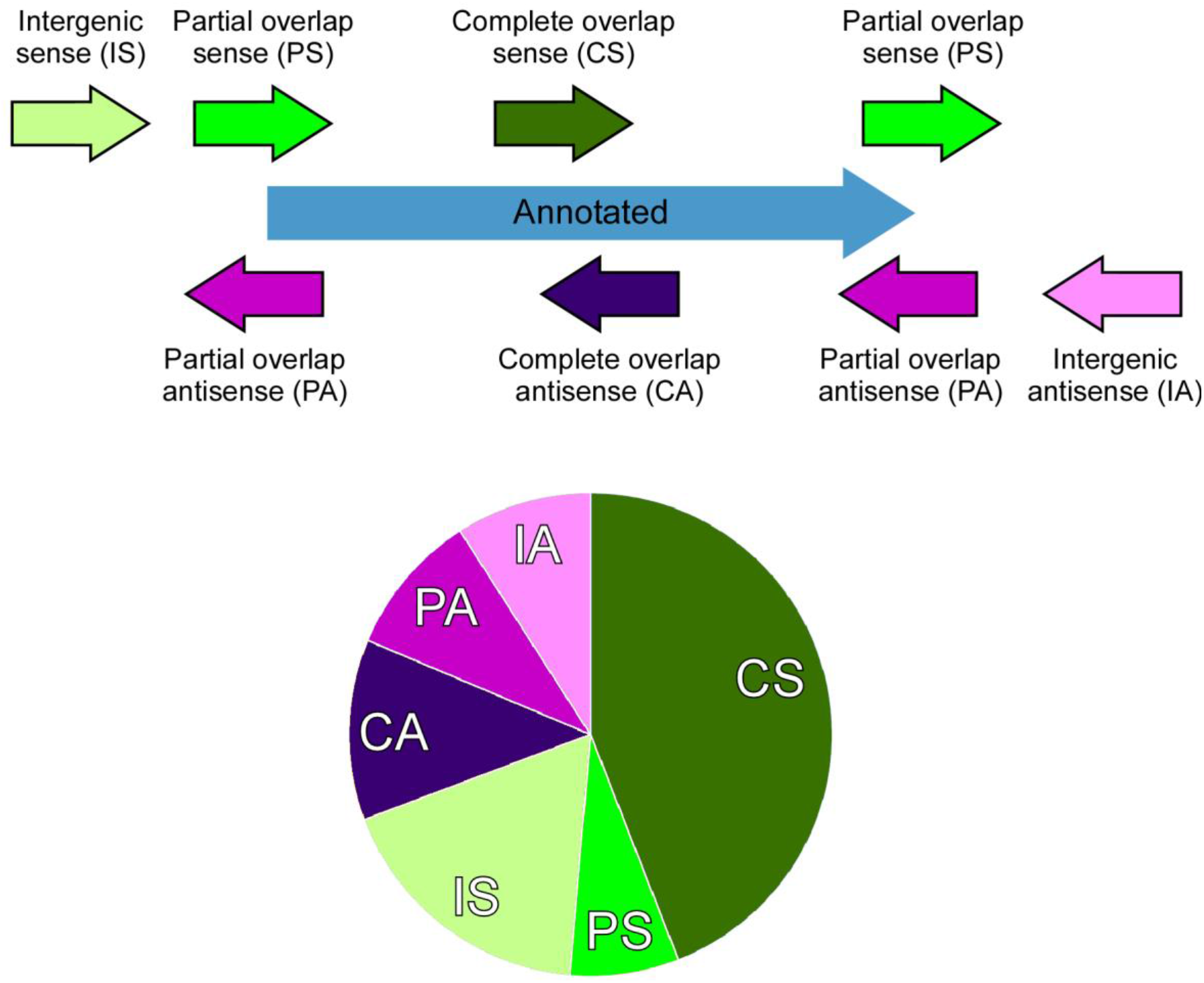
Classification of novel ORFs by position relative to annotated genes. The pie chart shows the distribution of novel ORFs across different categories defined by the type of overlap with an annotated gene, as shown in the schematic.

Strikingly, the sizes of novel ORFs are typically much smaller than those of annotated ORFs; the median length of novel ORFs is 15 codons, and 365 of the novel ORFs encode proteins of ≤50 amino acids, and hence are classified as sORFs. We compared the 365 putative novel sORFs found in *E.* coli BL21 with sORFs described for *E. coli* K-12 (strain MG1655); 15 of the putative sORFs perfectly match those described previously for MG1655, and 2 of the putative sORFs match previously described ORFs with one or two nucleotide mismatches (3, 14, 15).

### Experimental validation of putative novel sORFs

We selected 18 sORFs for experimental validation, covering a range of expression values based on Ribo-RET and Ribo-Api/Pmn coverage at start and stop codons, respectively. We C-terminally epitope tagged 17 of these ORFs at their native loci with SPA tags. We then used western blotting to detect stably expressed tagged proteins for cells grown in rich medium. As a positive control, we used a strain expressing SPA-tagged YhgP, an sORF that was previously detected by western blotting (3). We observed single bands for the positive control protein and for 9 of the 17 proteins encoded by putative sORFs. We observed no bands for an untagged strain grown under the same conditions (Figure 4A). We conclude that approximately half of the putative sORFs are stably expressed. We assigned new gene names to the ORFs for which a protein was detectable by western blot (Figure 4; Table S3).

**Figure 4.**
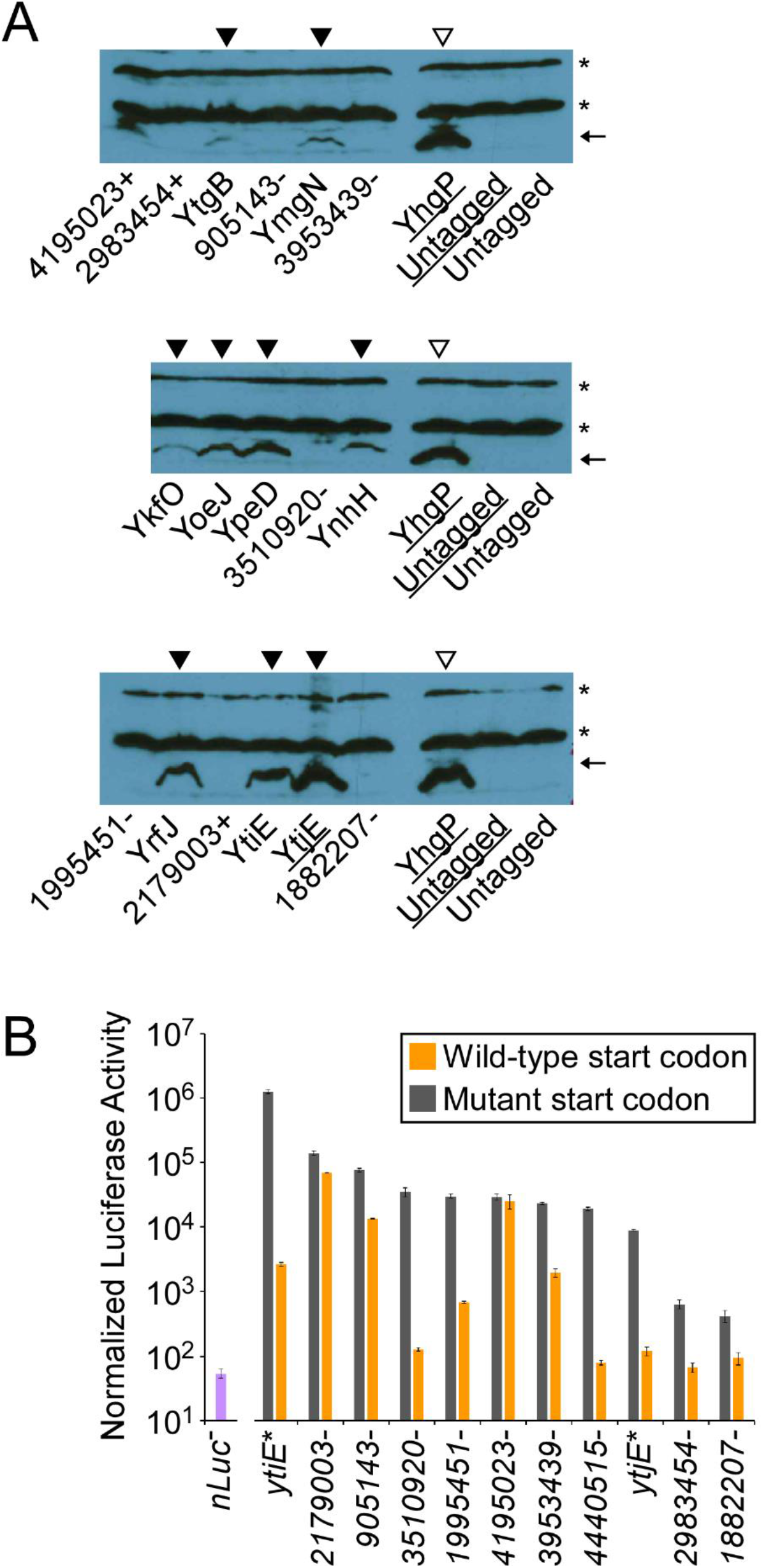
Experimental validation of sORFs by western blot and luciferase fusions. **(A)** Western blotting of SPA-tagged novel small proteins. Each lane was loaded with whole cell extract from cells expressing a SPA-tagged small protein, except for lanes labeled “Untagged”, which were loaded with wild-type MG1655. Underlined sample names indicate that the cells were Δ*thyA* and grown in medium supplemented with thymine. Asterisks indicate non-specific bands. The arrow indicates the expected position of SPA-tagged small proteins. YhgP is a positive control, previously confirmed by western blot. Untagged strains are negative controls. Empty triangles indicate the positive control lanes. Filled triangles indicate novel small proteins with bands of the expected size. **(B)** Luciferase assays for nano-luciferase (nLuc) fusions immediately downstream of the start codons of putative sORFs. Gray bars indicate normalized luciferase activity for nLuc fusions with wild-type start codons; orange bars indicate normalized luciferase activity for nLuc fusions with mutant (NCG) start codons. sORF names with asterisks indicate those for which we detected a corresponding protein by western blot. “nLuc^-^” (purple bar) indicates control data for cells without nLuc. Data represent the average of three independent biological replicates; error bars represent ± one standard deviation from the mean.

We reasoned that failure to detect a protein by western blot could either be due to the corresponding sORF being a false positive, or instability of the protein. To distinguish between these possibilities for the eight sORFs where the corresponding protein was not detected by western blot, for one sORF not tested by western blotting, and two of the sORFs where the corresponding protein was detected by western blot, we replaced the sORFs with a nano-luciferase gene at their native loci. We constructed equivalent strains with mutations in the putative start codons (ATG → ACG). For all ORFs tested, we detected robust luciferase expression; in all but one case, luciferase activity was significantly lower in the start codon mutant strain (Figure 4B). We conclude that these sORFs are actively translated, but the encoded proteins are not easily detected by western blot, likely due to instability. In some cases, mutation of the start codon did not reduce luciferase activity to background levels, suggesting that other in-frame trinucleotide sequences upstream of the predicted start codon can serve as alternative start codons. In the case of sORF 4195023- (named based on the genome coordinate and strand of the start codon), luciferase activity was the same for both wild-type and mutant constructs, suggesting that the major start codon is upstream of the predicted start codon.

### Evidence for function of novel sORFs

Our data strongly suggest that the large majority of the 397 novel sORFs are actively translated in *E. coli* BL21, but this does not indicate whether the sORFs or their encoded proteins are functional, with “function” defined as providing a positive contribution to cell fitness (22). Indeed, recent studies suggest that bacteria and eukaryotes can express hundreds of sORFs that are likely non-functional (4, 23, 24). We used three approaches to identify likely functional sORFs. First, we searched the encoded proteins for functional domains in the PFAM database (25). Most of the proteins are too short to include a functional domain; nonetheless, 6 of the novel ORFs have significant matches to functional domains listed in the PFAM database (Table S3), with the shortest of these ORFs encoding a protein of 61 amino acids. Second, we examined the codon bias of the sORFs; codon sequences that are non-random compared to the overall nucleotide content of the ORFs are indicative of purifying selection, a feature observed for annotated ORFs. We reasoned that novel sORFs with codon usage similar to annotated ORFs are likely under purifying selection. We analyzed the codon usage of 105 novel sORFs in regions that have at least 30 nt non-overlapping with annotated genes (including annotated genes not identified by Ribo-RET or Ribo-Api/Pmn). Specifically, we calculated the Relative Codon Deoptimization Index (RCDI), a measure of how closely the codon usage of an ORF matches that of the annotated ORFs from the same genome (26, 27). 11 of the 105 selected sORFs have codon usage similar to that of annotated genes (RCDI score <1.9, *p* < 0.01; RCDI scores closer to 1 indicate greater codon optimality; see Methods; Table S3), suggesting that these sORFs are under purifying selection, and hence are likely to be functional. Third, we searched for homologues of proteins encoded by novel sORFs, excluding those that have previously been described in *E. coli* MG1655, and limiting the analysis to proteins encoded entirely in intergenic regions. Thus, we identified likely homologues for seven of the encoded proteins (Table S3; Figure 5). Overall, our data identify a subset of sORFs that are likely to be functional; however, we found no evidence of function for the majority of sORFs.

**Figure 5.**
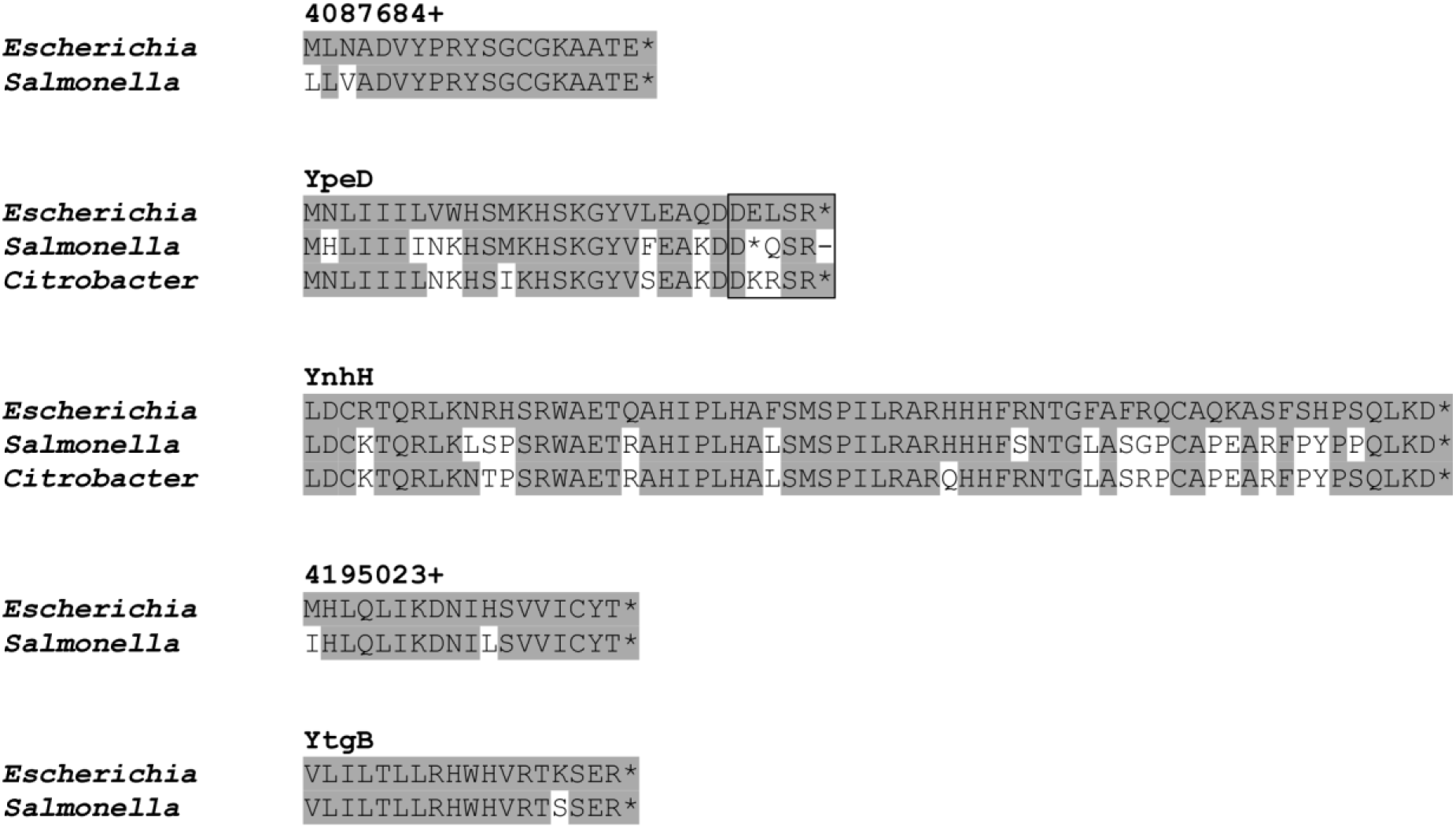
Conservation of selected sORFs. Sequence alignments of five selected novel proteins and their putative homologues in *Salmonella enterica* and (where found) *Citrobacter koseri*. Alignments were extracted from tBLASTn output files. Shaded amino acids match the protein sequence from *E. coli*. The boxed region for YpeD and putative homologues indicates the region of overlap in the corresponding genes with the downstream gene (*mntH*) and its homologues. Representative *S. enterica* and *C. koseri* genome sequences are shown (*Salmonella enterica* subsp. enterica serovar Worthington, strain CFSAN051295, CP029041.1; *Citrobacter koseri*, strain FDAARGOS_1029, CP066089.1).

## DISCUSSION

### Combining Ribo-RET and Ribo-Api datasets is a highly specific approach to identify translated ORFs

Previous studies in prokaryotes and eukaryotes have identified start codons by combining Ribosome Profiling coupled with antibiotics such as retapamulin or homoharringtonin that trap initiating ribosomes (2, 3, 28–31). While this approach has been effective at mapping sites of translation initiation, addition of the antibiotics does not completely remove signal from elongating ribosomes. Hence, Ribo-RET and related approaches have uncertainty that can make it difficult to identify start codons with high confidence, especially for start codons internal to highly expressed ORFs (29). Moreover, Ribo-RET signal at a putative start codon does not indicate whether ribosomes actively translate the corresponding ORF to completion in the absence of antibiotic.

Similar to Ribo-RET and related approaches, Ribo-Api/Pmn can be used to identify sites of translation termination, i.e. active stop codons. However, apidaecin treatment does not cause ribosomes to be exclusively associated with stop codons; ribosomes are also enriched in the regions immediately upstream and downstream of stop codons, and at start codons (19), such that stop codon identification from Ribo-Api/Pmn data is associated with a high FDR (estimated at 33.7% for our analysis). Moreover, identification of a stop codon position rarely indicates the position of the associated start codon(s).

Since IERFs in Ribo-RET data and TERFs in Ribo-Api/Pmn data are independent measurements, but start codons can be unambiguously associated with a single stop codon, combining data from both approaches allows ORF identification with a greatly reduced FDR compared to Ribo-RET or Ribo-Api/Pmn alone (estimated at 2.2% for our analysis). Moreover, detecting ribosomes at both start and stop codons for the same ORF strongly suggests that translation runs from start to stop codon.

The high sensitivity of an approach combining Ribo-RET and Ribo-Api/Pmn is evidenced by the fact that 35% of novel ORFs and 9% of annotated ORFs we identified have only two Ribo-Api/Pmn sequence reads at their stop codons. Nonetheless, the low FDR, and the propensity for novel ORFs to have low RNA secondary structure around their start codons strongly suggest that the majority of novel ORFs are actively translated. We anticipate that Ribo-RET and Ribo-Api/Pmn or related approaches will be combined to identify translated ORFs with high confidence in other bacterial and non-bacterial species. One potential barrier to these methods is the ability of the drugs to accumulate within cells; this could be prevented due to low permeability, or active efflux. While Ribo-RET has proved effective in *E. coli* (2) and *M. tuberculosis* (4), deletion of the *tolC* gene encoding a component of the drug efflux pump was needed for successful Ribo-RET with *E. coli*, and most Gram-negative species have similarly low sensitivity to retapamulin. Apidaecin uptake requires the *sbmA* gene, which encodes an inner-membrane transport protein (32); *sbmA* is missing in some Gram-negative and all Gram-positive bacterial species. If apidaecin uptake is low, an alternative approach would be to encode and express apidaecin, a small protein, inside the bacteria (33, 34). Similarly, Onc112, which functions similarly to retapamulin, is a small protein that could be encoded and expressed within cells (35).

### E. coli expresses large numbers of small proteins, many of which are likely unstable

Most of the novel ORFs we identified in this study are sORFs, i.e. they encode proteins of <51 amino acids. sORFs and small proteins have been challenging to identify by other methods; thus, we have greatly expanded the number of novel ORFs known for *E. coli*, with the caveat that not all have been experimentally validated using an independent approach. Luciferase assays measuring the capacity of the translation initiation region of selected novel ORFs at their native loci to direct translation indicate that expression levels vary greatly between sORFs, although it is important to note that the native sequences immediately downstream of the start codons of these sORFs were replaced in these assays by introduction of the nLuc gene; differences in expression caused by these downstream sequences will have been lost. Western blotting also indicates large differences in expression of small proteins encoded by selected novel sORFs. While differences in protein levels detected by western blot roughly mirrored differential expression levels measured using luciferase reporters, there were notable differences. In particular, protein levels of YtjE were moderately higher than those of YtiE, whereas luciferase activity for the *ytiE-nLuc* reporter was >100-fold higher than that for the *ytjE-nLuc* reporter. These differences are likely attributable to differences in stability between small proteins.

Nine of the proteins encoded by selected sORFs were not detectable by western blot, yet we detected robust luciferase activity for all the sORF-*nLuc* fusions. These data strongly suggest that (i) the large majority of novel sORFs identified by combining Ribo-RET and Ribo-Api/Pmn are actively translated, and (ii) many of the proteins encoded by identified sORFs are unstable.

### *Evidence that most small ORFs in* E. coli *are not functional*

Several lines of evidence suggest that most of the *E. coli* sORFs we identified are not functional, with “function” defined as providing a positive contribution to cell fitness, and hence being under selective pressure (22): (i) the codon usage of most analyzed novel sORFs did not match that of annotated ORFs; (ii) we detected sequence conservation at the amino acid level for only a few of the novel sORFs in genera beyond *Escherichia* and *Shigella* (*Shigella* species were excluded due to very high similarity between *Shigella* and *Escherichia* sequences). These analyses, however, have important limitations: (i) codon usage patterns are only informative for sORFs that do not overlap annotated ORFs; (ii) the statistical significance of both codon usage and sequence conservation is limited by the small size of sORFs; (iii) regulatory sORFs can be functional even if the corresponding proteins are not, and these sORFs often have non-standard codon usage. Based on our study of sORFs in *M. tuberculosis*, we proposed that the *M. tuberculosis* transcriptome is pervasively translated, with most translated proteins being non-functional. We propose that the *E. coli* is also subject to pervasive translation, although seemingly at a level below that of *M. tuberculosis* based on the relative number of novel and annotated ORFs identified in the two studies. A lack of function for most sORFs is consistent with half of the small proteins tested appearing to be weakly expressed and/or unstable, features expected of spurious proteins.

### Annotated genes that contain start codons for novel ORFs tend to have suboptimal codon usage

The position of novel ORFs with respect to annotated genes is non-random (Figure 3). Specifically, we detected many more novel ORFs on the same strand as annotated overlapping or downstream genes. We speculate that this bias is because novel ORFs require a transcript, and mRNAs for annotated genes tend to be more abundant than antisense RNAs. Nonetheless, we were surprised to find that 44% of all the novel ORFs we detected overlap completely with an annotated gene on the same strand; this suggests that overlapping translation of the annotated ORFs does not always prevent translation initiation at ORF-internal start codons, as suggested previously (2).

We speculated that novel ORFs whose start codons are within annotated genes reflect inefficient translation of the overlapping annotated ORF. To test this idea, we assessed the codon usage of annotated genes that encompass novel ORF start codons. The codon usage of annotated genes that encompass novel ORFs (mean RCDI score = 1.41) is significantly less optimal than that for the set of all other annotated ORFs (mean RCDI score = 1.33; Mann-Whitney U-test comparing RCDI scores, *p* = 1.7e^-5^). This difference was more pronounced when we considered only highly expressed novel ORFs (mean RCDI score of 1.59 for annotated genes that encompass novel ORF starts with Ribo-RET signal >5 RPM; *p* = 2.1e^-12^). We conclude that suboptimal codon usage of an annotated ORF increases the likelihood of internal initiation sites, likely due to a reduced efficiency of translation elongation. Suboptimal codon usage is a feature of horizontally acquired genes in *E. coli* (36), suggesting that horizontally acquired genes often have an increased abundance of internal translation. Another feature of horizontally acquired genes is low G/C content (36). Strikingly, annotated genes that encompass highly expressed novel ORF starts have a much lower G/C content (mean of 42.2%) than the set of all annotated ORFs (mean of 51.1%). Given that genes with internal start codons often have suboptimal codon usage, and often have low G/C content, we speculate that horizontally acquired genes are hotspots for internal novel ORFs due to inefficient translation elongation.

### *Identification of* E. coli *sORFs that are likely to be functional*

A few of the novel sORFs we identified are associated with features expected of functional proteins. Specifically, we detected stably expressed protein by western blot, we observed codon usage patterns similar to those of annotated ORFs, and/or we identified putative homologues in other genera. While the functions of these sORFs or their corresponding small proteins are unknown, they represent promising candidates for further study. In the case of sORFs that are immediately upstream or partially overlapping with annotated genes on the same strand, a potential function is regulation of the downstream gene by translation the sORF, as has been described for other sORFs in bacteria (17). Our data also suggest exercising caution when selecting sORFs/small proteins for further study, since many sORFs/small proteins may be non-functional; relying on codon usage patterns and/or sequence conservation is a simple way to prioritize likely functional sORFs/small proteins.

## MATERIALS AND METHODS

### Strains and plasmids

All strains and plasmids used in this work are listed in Table 1. All oligonucleotides used in this work are listed in Table S4. All strains used in this work are derivatives of *Escherichia coli* K-12 MG1655 (37). Strains AMD783-AMD800 were constructed using λ Red recombineering as described previously (38). Specifically, PCR products used for recombineering were generated with oligonucleotides JW10796-JW10803 and JW10883-JW10912, using DY330 *allR-SPA::kanR* (39) as a template. The PCR products generated to make AMD783 and AMD784 were electroporated into AMD052 (*Escherichia coli* K-12 MG1655 Δ*thyA*) harboring pKD46 (38). The PCR products generated to make AMD785-AMD801 were electroporated into MG1655 harboring pKD46.

**Table 1.**
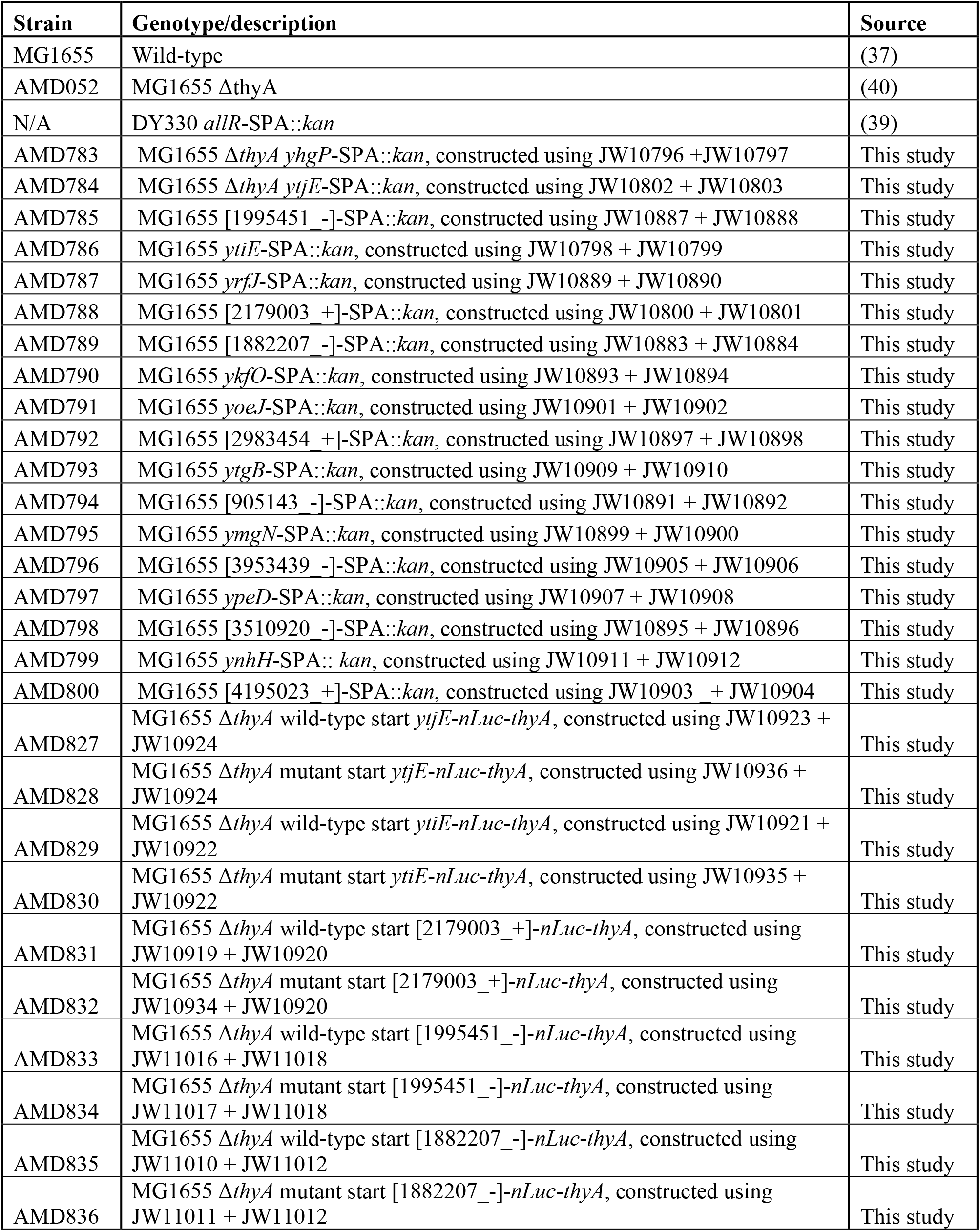
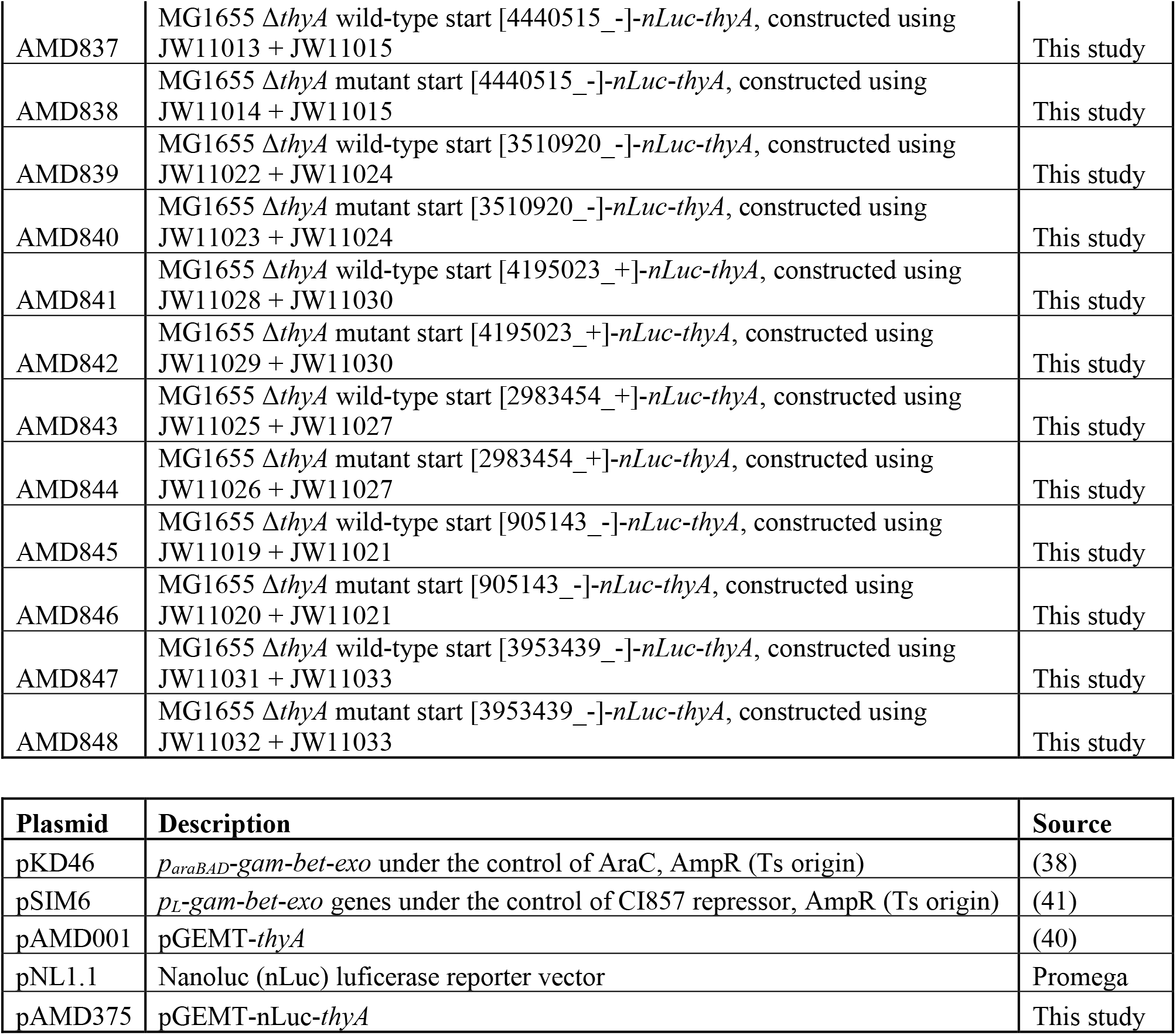
List of strains and plasmids used in this study.

Strains AMD827-AMD848 were constructed using the previously described FRUIT method of recombineering (40). Oligonucleotides JW10919-JW10924, JW10934-JW10936 and JW11010-JW11033 were used to generate PCR products for recombineering using pAMD375 (see below) as a template. The PCR products generated were electroporated into AMD052 harboring pSIM6 (41).

Plasmid pAMD375 was constructed using oligonucleotides JW10881 and JW10882 to amplify the NanoLuc luciferase gene from pNL1.1 (Promega). The resulting DNA fragment was cloned into pAMD001 (40) digested with *Apa*I and *Nco*I, using the In-Fusion cloning system (Takara).

### Identification of IEFRs from Ribo-RET data

Sequence reads for the Ribo-RET data (NCBI SRA accession number SRR8156054) (2) were aligned to the *E. coli* BL21 reference genome and to a reverse complemented reference genome using Rockhopper (version 2.03). The genome positions and strands of sequence read 3’ ends were inferred from the resultant .sam file. Sequence read coverage across known non-coding RNAs (Table S5) was set to zero, and coverage was normalized to reads per million (RPM). Every genome position on both strands was considered as a possible IEFR. IEFRs were selected if the corresponding position/strand met three criteria: (i) normalized sequence read coverage was ≥0.5 RPM; (ii) normalized sequence read coverage was ≥10-fold higher than the mean sequence read depth at all positions in a 101 nt window centered on the position being considered; (iii) normalized sequence read depth was at least as high as that for any other position in a 21 nt window centered on the position being considered.

### Identification of putative start codons from IEFRs

The frequency of all trinucleotide sequences was determined for positions −50 to −1 nt relative to the position of each IEFR. We reasoned that trinucleotide sequences used as start codons would be enriched relative to the local background in the −14 to −18 nt range relative to IEFR positions, with the strongest enrichment at position −15 (2); we considered the local background frequency for each trinucleotide sequence to be that for positions −50 to −41 nt relative to the IEFRs. To be considered as a possible start codon, we first selected candidate start codon sequences by requiring that a trinucleotide sequence be enriched at least 1.4-fold above background at position −15 relative to IEFRs, and that the highest level of enrichment in the −14 to −18 nt range be at position −15. These criteria were met by only three trinucleotide sequences: ATG, GTG, and TTG; hence, only these sequences were considered as possible start codons. ATG is enriched at least 1.4-fold above background at positions −14 to −18 nt relative to IEFRs; GTG is enriched at least 1.4-fold above background at positions −14 to −16 nt relative to IEFRs; TTG is enriched at least 1.4-fold above background at positions −14 to −15 nt relative to IEFRs. We therefore identified putative start codons as ATG sequences positioned 14-18 nt upstream of IEFRs, GTG sequences positioned 14-16 nt upstream of IEFRs, and TTG sequences positioned 15-16 nt upstream of IEFRs.

### Calculating the False Discovery Rate (FDR) for putative start codons

The likelihood of randomly selecting a genome coordinate with an associated start codon sequence (as defined above for IERFs) was estimated by selecting 100,000 random genome coordinates and determining the fraction, “R”, that would be associated with a start codon. The set of IERFs contains a number of true positives (i.e. corresponding to a genuine start codon), and a number of false positives. We assume that true positive IERFs are all associated with a start codon using the parameters described above for calling ORFs. We assume that false positive IERFs are associated with a start codon at the same frequency as random genome coordinates, i.e. R. Since we know how many IERFs were not associated with a start codon, we can use this number to estimate how many false positive IERFs were associated with a start codon by chance. With the total number of IERFs as “I” and the total number of identified ORFs as “O”, the FDR for ORF calls is estimated by:

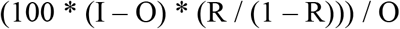

### Identification of TERFs from Ribo-Api/Pmn data

TERFs were identified as described for IEFRs, but using Ribo-Api/Pmn data (NCBI SRA accession number SRR11728142) (19).

### Control Ribo-seq data for puromycin-treated cells

Sequence read coverage for a control dataset (puromycin-treated cells) were generated as described above (NCBI SRA accession number SRR11728143) (19).

### Identification of putative stop codons from TERFs

Putative stop codons were identified from TEFRs using the same approach as described above for identifying putative start codons from IEFRs, except that only TAA, TGA and TAG trinucleotide sequences were considered. Based on the enrichment of TAA, TGA and TAG sequences above the local background in the regions upstream of TEFRs, we identified putative stop codons as TAA, TGA or TAG sequences positioned 12-14 nt upstream of TEFRs. The FDR for putative stop codons was determined as described for start codons.

### Combining IERFs and TERFs to identify putative ORFs

Stop codons identified from TERFs, were discarded if the TERF was located within 3 nt of an IERF on the same strand, since apidaecin stalls ribosomes at start codons in addition to stalling ribosomes at stop codons. Putative start codons identified from IERFs were translated *in silico* to identify the corresponding stop codon. These stop codons were compared to stop codons identified from TERFs; for shared stop codons, the ORF defined by the IERF was selected as a putative ORF.

To conservatively estimate an FDR for ORF identification, we rotated the normalized sequence read coverage data for Ribo-RET and Ribo-Api/Pmn relative to the genome sequence by 1,000 nt, each of 10 times. After each cycle of rotation, we repeated IERF, TERF, and ORF identification, as described above. The average number of ORFs identified after each of the 10 rotations was considered to be a conservative estimate of the number of falsely discovered ORFs in the non-rotated data. FDRs for annotated, isoform and novel ORFs were estimated by determining how many of the ORFs identified from rotated data were found in each category, and dividing the overall FDR equivalently.

### RNA folding prediction

The sequence from −25 to +15 nt relative to each start codon, or for 500 x 41 nt sequences randomly selected from the *E. coli* BL21 genome, were selected for prediction of the free energy of the predicted minimum free energy structure with a local installation of ViennaRNA Package tool RNAfold, version 2.4.14, using default settings (42).

### Western blotting

MG1655 (untagged) and strains AMD785-AMD800 were grown overnight at 37 °C in LB, subcultured 1:100 in LB, and grown with shaking at 37 °C to an OD_600_ of 0.6–0.7. AMD052, AMD783 and AMD784 were grown overnight at 37 °C in LB supplemented with 100 μg/mL thymine, subcultured 1:100 in LB supplemented with 100 μg/mL thymine, and grown with shaking at 37°C to an OD_600_ of 0.6–0.7. After growth, 500 μL of each culture was pelleted and resuspended in 25 μL Laemmli sample buffer with a 19:1 ratio of buffer: β-mercaptoethanol. Samples were separated on a 10% SDS-PAGE gel. Proteins were transferred to PVDF membranes and probed with M2 anti-FLAG antibody (Sigma; 1 in 5,000 dilution) and HRP-conjugated goat anti-mouse antibody (1 in 250,000 dilution). Tagged proteins were visualized using the SuperSignal™ West Femto detection kit (ThermoFisher).

### Luciferase reporter assays

Strains AMD825-AMD848 were grown overnight in LB at 37°C, subcultured 1:100 in LB and grown with shaking at 37°C to an OD_600_ of 0.4-0.7. After recording the OD_600_, 10 μL of each culture was aliquoted into a 96-well opaque plate with three technical replicates each. All aliquots were then mixed with 10 μL Nano-Glo (Promega). Luminescence readings (relative light units; RLUs) were taken using a Turner Biosystems Veritas microplate luminometer. Relative luminescence values were reported as RLU/OD_600_.

### Analysis of conserved protein domains

A FASTA file containing all proteins encoded by novel ORFs with >20 codons was submitted to the PFAM webserver (version 34.0) (25) for a batch sequence search using default parameters.

### Codon usage analysis

FASTA files containing the nucleotide sequences of regions of novel ORFs not overlapping with annotated genes were generated and submitted to the RCDI/eRCDI web server (27). The *E. coli* BL21 genome codon usage table (https://www.kazusa.or.jp/codon) was used, and the following parameters were used: the upper confidence limit was set to 99% confidence/99% population, the length of random sequence of 300 nt was selected, and the standard genetic code was used.

### Sequence conservation analysis

The amino acid sequences of all proteins encoded by novel, fully intergenic ORFs were submitted to the NCBI tBLASTn webserver (version 2.11.0) (43). The *Escherichia* (taxid:561) and *Shigella* (taxid:620) genera were excluded from the search. The following parameters were adjusted: 250 max target sequences were selected, the expect threshold was set to 100, and the low complexity region filter was unselected. The search results were further filtered by selecting only ORFs with 100% query coverage and a ≥70% sequence identity. Groups of conserved ORFs were realigned using the Clustal Omega multiple sequence alignment program with default settings (44).

## ACKNOWLEDGEMENTS

We thank the Wadsworth Center Applied Genomic Technologies Core Facility for DNA sequencing. We thank the Wadsworth Center Tissue Culture and Media Core Facility and Glassware Facility for technical support. We thank Alexander Mankin, Nora Vazquez-Laslop, and Gisela Storz for helpful suggestions, and comments on the manuscript. We thank Don Court for the recombineering plasmid pSIM6. KM was supported by a National Institutes of Health Training Grant (5T32AT007533). JTW was supported by National Institutes of Health grant R01GM139277.

## SUPPLEMENTARY TABLES

Table S1. List of IERFs.

Table S2. List of TERFs.

Table S3. List of ORFs identified by combining Ribo-RET and Ribo-Api/Pmn.

Table S4. List of oligonucleotides used in this study.

Table S5. List of genomic regions containing non-coding genes that were excluded from the Ribo-RET and Ribo-Api/Pmn analyses.

